# Combining membrane potential and calcium imaging in brain slices using the voltage sensitive dye ElectroFluor630 and the calcium indicator Calbryte520

**DOI:** 10.64898/2025.12.02.691797

**Authors:** Shirin Ghasemiform, Marco Canepari

## Abstract

Wide-field imaging from brain slices stained with a voltage sensitive dye (VSD) and simultaneously loaded with a Ca^2+^ indicator allows investigating neuronal excitability and synaptic transmission at multi-cellular scale. So far, achieving this type of combined imaging has been limited by experimental constraints. We assessed the ability of the red-IR emitting VSD ElectroFluor630 (EF-630) to be combined with blue-excitable green-emitting Ca^2+^ indicators to record signals elicited by electrical stimulation in hippocampal slices. Transversal mouse hippocampal slices were stained with EF-630. Ca^2+^ indicators, either Fluo-4, Fluo-8, Cal520 or Calbryte520, were loaded using their AM-ester forms. Fluorescence, during stimulation of the CA3 region was imaged at 5 kHz from hippocampal areas of ∼750X250 µm^2^ at 1 µm pixel resolution. After assessing all Ca^2+^ indicators, we selected Calbryte520 for achieving >30 minutes stable recordings in combination with EF-630. Action potentials and related Ca^2+^ transients were detected in the CA3 stimulated area whereas synaptic signals were observed in the CA1 region. On these signals, we tested the pharmacological blockade of either action potentials or glutamatergic synaptic potentials. We report novel optical measurements of both electrical and Ca^2+^ transients in brain slices, providing unique information on neuronal excitability and network activity.

## 1 Introduction

Membrane potential (V_m_) imaging in brain slices using a voltage sensitive dye (VSD)^1,2^ allows monitoring electrical neuronal activity from small populations of cells, enabling the investigation of both neuronal excitability and synaptic transmission. Separately, Ca^2+^ imaging in brain slices loaded with AM-ester dyes^3,4^ allows measuring transients of intracellular Ca^2+^ concentrations that can be related, in different manners, to the electrical activity. Thus, complementary information obtained by combining the two optical recordings might permit deep understanding of network activity. V_m_ imaging has been performed in slices from several brain areas including the cortex^5,6,7,8,9,10,11^, the hippocampus^12,13,14,15,16^, the olfactory bulb^17^ and the cerebellum^18^, with various absorbance or fluorescence VSDs^19^ including di-4-ANEPPS, RH795, RH414, RH479, RH155, RH482. Using these dyes, slices were stained for several minutes and fluorescence changes (ΔF/F_0_) associated with V_m_ transients were in the range of 0.05-0.5%, requiring collection of large numbers of photons to be distinguished from photon “shot” noise. This was normally achieved by using fast photodiode arrays, at the cost of lowering spatial resolution, and only recently by using CMOS cameras with higher spatial resolution^20,21^. Regarding Ca^2+^ imaging in brain slices, this was done using several high-affinity indicators, in particular probes with fluorescence excited by blue wavelengths and emitting green wavelengths (like “fluorescein”), such as Calcium Green^22^, Oregon Green BAPTA-1^23^, Fluo3^24,25^ or Fura4^26,27,28^. Ca^2+^ activity was recorded after incubation of brain slices with the AM-ester forms of Ca^2+^ indicators normally for >30 minutes. In principle, V_m_ and Ca^2+^ imaging can be combined in the same preparation if fluorescence spectra of the two dyes are distinct. In practice, the ability of V_m_ and Ca^2+^ imaging to explore neuronal excitability and synaptic transmission, as well as to combine the two techniques, is constrained by several technical aspects which are listed below.

1. The efficiency of both VSDs and Ca^2+^ indicators to stain (or load) neurons must be adequate, and the amplitude of the resulting signals (ΔF/F_0_) must be large enough, to produce measurements with sufficient signal-to-noise ratio (SNR). Optical signal quality thus depends on the solubility of VSDs in water and on the penetration of AM-ester forms into cells, as well as on the efficiency of esterases to cleave the AM group releasing the indicator into the cytosol.
2. Biological experiments require dye stability to enable recordings for long periods. This is given by the VSD retention in the plasma membrane, or Ca^2+^ dye retention in the cytoplasm, as well as by the dye resistance to photodamage produced by light exposure.
3. For combination of V_m_ and Ca^2+^ imaging, ideally, there should be absolutely no emission of each dye in the wavelength recording window of the other one. In practice, the VSD RH414 was combined with Calcium Orange^29,30^ and the VSD Di-2-ANEPEQ with Calcium Green-1^31^, but in both combinations dyes had overlapping excitation. Thus the ability of obtaining “clean” V_m_ and Ca^2+^ signals relied entirely on each dye giving only a negligible component in the wavelength recording window of the other one. In studies with dyes injected into single cells, Di-2-ANEPEQ (JPW1114) and Oregon Green BAPTA5N had negligible crosstalk in recordings from CA1 hippocampal pyramidal neuron dendrites^32,33^, but not in dendrites of cerebellar Purkinje neurons^34^ and of layer-5 neocortical pyramidal neurons^35^. Thus, no-overlapping excitation is also a necessary condition to always obtain independent V_m_ and Ca^2+^ recordings.

Here we report combined V_m_ and Ca^2+^ imaging measurements that extend this type of measurements from the single-cell approach, where dyes are injected into single cells, to a population imaging approach where brain slices are stained/loaded with VSD and Ca^2+^ indicators. V_m_ imaging was performed with the VSD Di-4-ANEQ(F)PTEA^36^, or ElectroFluor630 (EF-630), that was used in cardiac preparations^37^ and only recently in brain slices^38^. This indicator is soluble in water and can stain slices with ∼1 minute of exposure at micromolar concentrations. By stimulating with an extracellular electrode, the synchronised action potential (AP) in the adjacent area produced an absolute fluorescence transient >1% when fluorescence was excited at 640 nm. The slower V_m_ changes in other areas, associated with excitatory synaptic potentials (EPSPs) and asynchronous firing, could be as large as 0.5%. We assessed that these signals were stable for at least 30 minutes. Since EF-630 was excited at 640 nm and emits at longer wavelengths, this VSD can be combined with common blue-excited and green-emitting Ca^2+^ indicators. Among these dyes, we assessed the performance of Fluo4, Fluo8, Cal520 and Calbryte520 and established that the last one provides large (ΔF/F_0_ >1%) signals with stability comparable to that of EF-630. The relatively high amplitude of both V_m_ and Ca^2+^ signals allowed resolving them by acquiring images of ∼750X250 pixels at 5 kHz using a state-of-the-art CMOS camera. Thus, in a standard hippocampal preparation, we report an initial characterisation of functional imaging signals obtained with the unprecedented combination of EF-630 and Calbryte520.

## 2 Materials and Methods

### 2.1 Ethical approval

Experiments were performed at the Laboratory of Interdisciplinary Physics in Grenoble in accordance with European Directives 2010/63/UE on the care, welfare and treatment of animals. Procedures were reviewed by the ethics committee affiliated to the animal facility of the university (E3842110001). Mice (C57BL/6j) were kept in the animal house and fed *ad libitum* and anesthetised by isoflurane inhalation before euthanasia.

### 2.2 Slice preparation and solutions

After brain removal, transversal (horizontal) hippocampal slices with 350 µm thickness were prepared from 3-6 postnatal weeks old mice using a Leica VT1200 (Leica, Wetzlar, Germany) as recently described^28,39^. The extracellular solution contained (in mM): 125 NaCl, 26 NaHCO_3_, 20 glucose, 3 KCl, 1 NaH_2_PO_4_ and 2 CaCl_2_ bubbled with 95% O_2_ and 5% CO_2_. After slicing, slices were pre-incubated for 45 minutes at 37°C and then maintained at room temperature. During experiments, slices were perfused in the recording chamber with extracellular solution at 32-34°C. In some experiments, the following chemicals were added to extracellular solution: Tetrodotoxin (TTX); 2,3-Dioxo-6-nitro-1,2,3,4-tetrahydrobenzo[*f*]quinoxaline-7-sulfonamide disodium salt (NBQX); D-(-)-2-Amino-5-phosphonopentanoic acid (D-AP5). All chemicals were from Hello Bio (Dunshaughlin, Republic of Ireland) and they were pre-dissolved in water before being diluted to the final concentration indicated in the Results.

### 2.3 Slice loading with Ca^2+^ indicators and staining with EF-630 VSD

To load Ca^2+^ indicators, slices were incubated at room temperature for 30-45 minutes with extracellular solution containing 1-2 µM of the AM-ester pre-dissolved in DMSO at 5 mM. Assessed Ca^2+^ indicators were Fluo-4, Fluo-8, Cal520 and Calbryte520 (AAT Bioquest, Pleasanton, CA). Staining with EF-630 (Potentiometric Probes, Farmington, CT) was done directly in the recording chamber with a procedure similar that developed in another laboratory^38^. In detail, continuous prefusion was stopped and the volume of the extracellular solution was reduced to ∼200 µL. 3 µL of a solution containing 500 µM EF-630 dissolved in H_2_0 was added and the slice was kept for ∼1 minute under ∼7.5 µM EF-630 before perfusion was restarted. In combined imaging experiments, Calbryte520 loading was done before EF-630 staining.

### 2.4 Optical arrangement, electrical stimulation and imaging

The setup, based on an Olympus BX51 microscope equipped with a 25X/1.05 NA objective (model XLPLN25XWMP2), was used in recent studies^28,39^. The diameter of the field of view of this objective is ∼720 µm. As previously described^28,39^, to image with a pixel resolution of ∼1 µm from the largest possible portion of the hippocampus, images projected to a Kinetix (Teledyne Photometrics, Tucson, AZ) CMOS camera, used for imaging, were de-magnified by 0.25X. In the mouse hippocampus, this arrangement allowed visualising with ∼750 pixels in the X-direction the last part of the CA3 pyramidal cell layer and most of CA1 pyramidal cell layer. Images with ∼250 pixels in the Y-direction were acquired at 5 kHz frame rate at 8-bit depth. For electrical stimulation, pipettes of 2-4 µM diameter filled with extracellular solution were used. Precisely, the pipette was positioned above the *stratum lucidum* corresponding to the last part of the mossy fibre pathway, i.e. of the CA3 region. Electrical activity was elicited by constant current pulses of 30-60 µA amplitude and 100 µs duration. EF-630 fluorescence was excited using the 640 nm line of an LDI-7 laser (89 North, Williston, VT) and emitted fluorescence was band-pass filtered at 728±64 nm (Fig 1a). Ca^2+^ fluorescence (from all indicators) was excited by a 470 nm OPTOLED (Cairn Research, Faversham, UK) and emitted fluorescence was band-pass filtered at 530 ± 21 nm (Fig 1b).

**Fig. 1.**
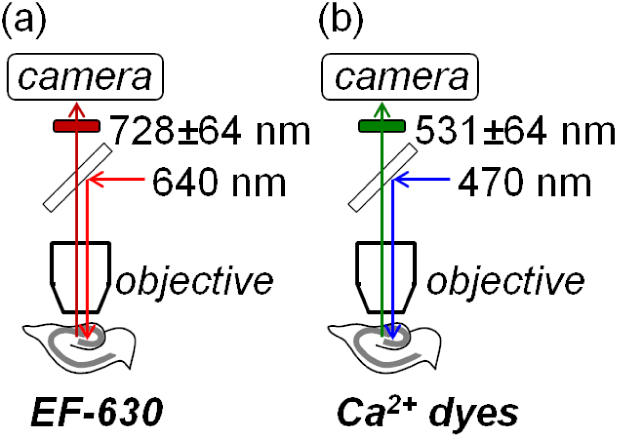
Scheme of the optical pathways used for the combination of two indicators. (a) For the VSD EF-630, fluorescence was excited at 640 nm and emitted light band-pass filtered at 728±64 nm before camera acquisition. (b) For Ca^2+^ indicators, fluorescence was excited at 470 nm and emitted light band-pass filtered at 530±21 nm before camera acquisition.

### 2.5 Data recording and analysis

Images were acquired with the open source software Micro-Manager and data were analysed either in MATLAB or in Python. Frame sequences, either from single trials or from averages of 2 or 7 trials with identical responses, were corrected for photo-bleaching. Signals were expressed as absolute ΔF/F_0_: +ΔF/F_0_ for Ca^2+^ fluorescence or -ΔF/F_0_ for V_m_ fluorescence since depolarisation corresponds to a decrease of fluorescence under our experimental conditions. In the whole article, they were calculated from regions-of-interest (ROI) of either 50X50 or 100X100 pixels. In one figure, signals were illustrated in coloured scales after applying twice a 50X50 median filter, first to the raw frames and then to the absolute ΔF/F_0_ frames. Finally, for quantification of slow signals beyond the noise, a Savitsky-Golay smoothing filter was utilised^40^.

### 2.5 Statistical analysis

In this technical report, paired t-tests were applied to assess the following hypotheses: (1) whether a rundown of fluorescence transients occurred; (2) whether parameters of signals differed at different sites in the same experiment; (3) whether delivery of a chemical had an effect on signal parameters. The test was applied to N = 5 slices when the effect of TTX was assessed, and to groups of N = 7 slices in all other cases. Values of P below certain thresholds (either 0.01 or 0.001) are reported in the text to support significant differences.

## 3 Results

### 3.1 EF-630 fluorescence transients following CA3 electrical stimulation

As described in the Materials and Methods, transversal slices of the hippocampus were loaded with the VSD EF-630 by exposing them to the dye in the recording chamber. After that, a portion of the pyramidal cell layer comprising the final part of the CA3 region and the initial part of the CA1 region was visualised (scheme on the top of Fig.2a). As in experiments presented in an earlier report^28^, a stimulating electrode was positioned above the *stratum lucidum* at the end of the mossy fibre (*MF*) pathway. A transmitted light image and an EF-630 fluorescence image after staining are shown in Fig.2a (bottom). At the same spatial resolution, fluorescence images could be acquired at 5k frames/s. In this example, five ROIs of 50X50 pixels (50X50 µm^2^) were selected, the first one (ROI*1*) centred on the tip of the stimulating electrodes and the others (ROIs *2-5*) in the *stratum radiatum* at progressively longer distance towards the CA1 region.

**Fig. 2.**
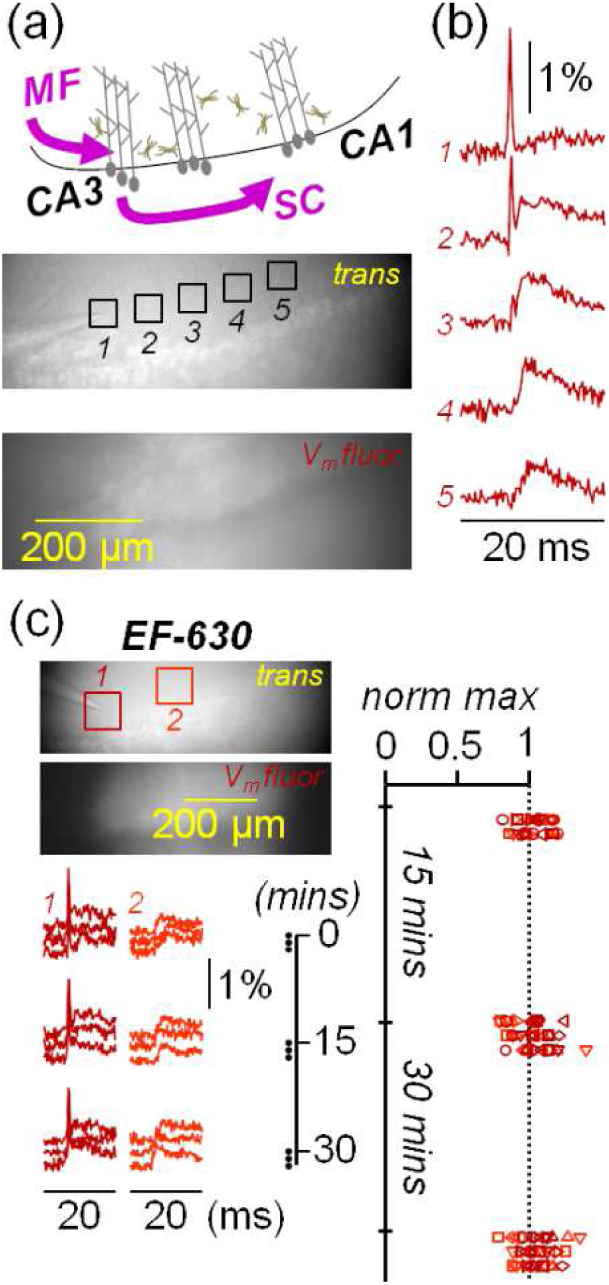
EF-630 transients in hippocampal slices. (a) Top, scheme of imaged hippocampal portions; the stimulation electrode is placed above the ending of *MF* pathway; the recording area includes most of the CA1 region where Shaffer collaterals (*SC*) form synaptic contacts with pyramidal neurons. Bottom, transmitted light (*trans*) image and EF-630 fluorescence (*V_m_ fluor*) image of the recording area (760X251 pixels) in a hippocampal slice; five ROIs of 50X50 pixels are indicated. (b) EF-630 (V_m_) transients (-ΔF/F_0_), from ROIs in panel a, associated with a stimulation pulse occurring at t∼8 ms in a 20 ms recording; a spike is detected in ROI1 and ROI2; depolarisations with EPSP shape are detected in ROIs 2-5. (c) Left-top, in another slice *trans* and *V_m_ fluor* images of the recording area; two 100X100 pixels ROIs are indicated. Left-bottom, nine consecutive V_m_ transients from the two ROIs above associated with a stimulation pulse; the first three recordings were initially performed every minute; the next three recordings were performed every minute after 15 minutes; the last three recordings were performed every minute 30 minutes after the first recording; spikes are detected in ROI1 (dark red) and EPSPs are detected in ROI2 (light red). Right, scatter plots of EF-630 transients maxima normalised to maxima of the first recording from N = 7 slices where the test of the example illustrated on the left was performed.

Fluorescence transients (averages of 7 trials) from the five ROIs are shown in Fig.2b. At ROI*1* and ROI*2*, a spike with a clear shape of an AP, presumably occurring in directly stimulated neuronal populations, is detected just after the stimulation pulse. At ROIs *2-5*, smaller depolarisation transients were observed. These signals have shape of excitatory postsynaptic potentials (EPSPs), but depolarisation in these neuronal populations can be boosted by asynchronous firing elicited by the EPSPs. In agreement with what reported by another laboratory^38^, we noticed that individual fluorescence transients were stable over time for many trials. Since we estimate that 30 minutes is a reasonable time window necessary for many types of studies in brain slices, we quantitatively assessed the stability of EF-630 over this time window. In the example of Fig.2c, fluorescence transients in single trials, from a 100X100 pixels (ROI (*1*) centred on the tip of the stimulator, and from another 100X100 pixels ROI (*2*) centred at ∼200 µm from the first one, are shown. We used larger ROIs in this case in order to quantify fluorescence transients from single trials. A stimulation pulse was delivered three times every minute, then three times after 15 minutes and finally three times after 30 minutes from the first stimulation. Transients from both ROI*1* (dark red) and ROI*2* (light red) were stable over time with fluctuations within the noise. We performed this test in N = 7 slices and obtaining the same result (scatter plots on Fig.2c). We concluded that EF-630 imaging can reliably monitor neuronal excitability and synaptic transmission in healthy brain slices for at least 30 minutes. Since the aim of this study was to characterise the ability of combining EF-630 imaging with Ca^2+^ imaging using blue-excitable green-emitting indicators, we assessed Ca^2+^ indicators in the same manner.

### 3.2 Assessment of four blue-excitable green-emitting Ca^2+^ indicators

We recently established the ability of the Ca^2+^ indicator Fluo4, previously used to monitor large slow signals from glial cells^26,27^, to report neuronal activity^28^. Using this dye, however, we observed a rundown of ΔF/F_0_ signals occurring within a few minutes. In the example of Fig. 3a, a pulse of stimulation was initially given three times every minute. Fluorescence transients were shown from a 100X100 pixels ROI, centred on the tip of the stimulator (dark green), and from another 100X100 pixels ROI, centred at ∼200 µm from the first one (light green). The first three signals recorded every minute were stable in both regions, but when we performed the next three recordings after 15 minutes signals were appreciably smaller. The same behaviour was observed in N = 7 slices where Fluo4 was tested (see scatter plots on Fig.3a) and the mean of the three signals recorded 15 minutes after the first trial were significantly smaller with respect to the first signal (P < 0.001, paired t-test). Since Fluo4 cannot perform over time as EF-630, we next assessed the similar indicator Fluo8. As shown in the example of Fig.3b and in the scatter plots on the right, reporting the measurement in N = 7 slices, Fluo8 signals are affected by rundown similar to that observed in slices loaded with Fluo4. Also for Fluo8, signals recorded 15 minutes after the first trial were significantly smaller than the first signal (P < 0.001, paired t-test).

**Fig. 3.**
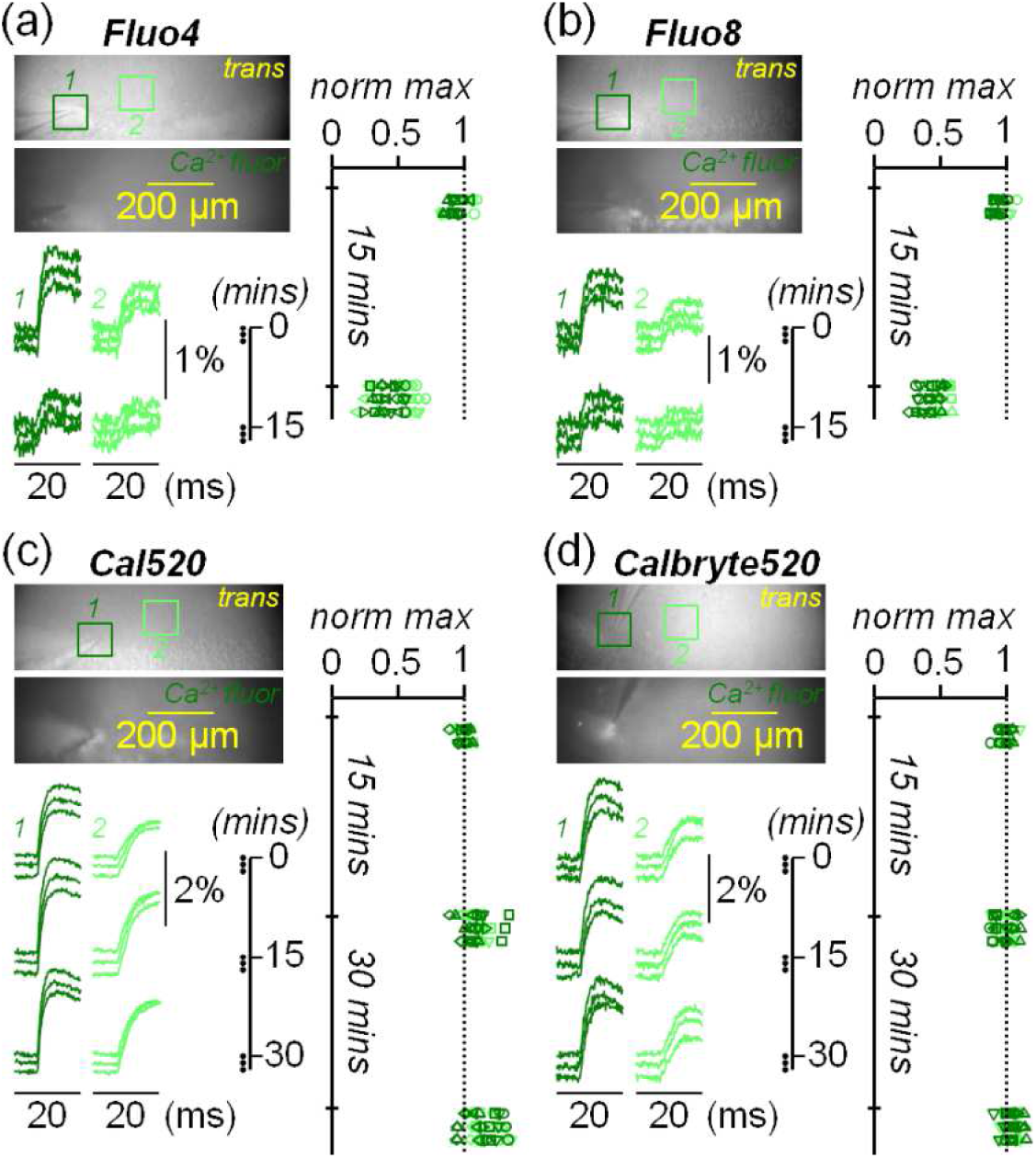
Ca^2+^ transients from four different indicators in hippocampal slices. (a) Left-top, in a slice loaded with Fluo4, *trans* and *Ca^2+^ fluor* images of the recording area; two 100X100 pixels ROIs are indicated. Left-bottom, six consecutive Ca^2+^ transients from the two ROIs above associated with a stimulation pulse; the first three recordings were initially performed every minute; the next three recordings were performed every minute after 15 minutes. Right, scatter plots of EF-630 transients maxima normalised to maxima of the first recording from N = 7 slices where the test of the example illustrated on the left was performed; a rundown of signals after 15 minutes is observed. (b) Same as a, but for slices loaded with Fluo8. (c) Same as a and b, for slices loaded with Cal520 but in this case with three additional recordings 30 minutes after the first recording; no rundown is observed in scatter plots on the right. (c) Same as c, for slices loaded with Calbryte520; also with this indicator, no rundown is observed in scatter plots.

In a comparative study where several Ca^2+^ indicators were tested^41^, it was established that Cal520^42^ performs better than Fluo4 and Fluo8 and we therefore assessed this dye. As shown in the example of Fig.3c, Cal520 gives transients that are larger than those from the previous two indicators, but more importantly these transients did not exhibit rundown when the three recordings were repeated first after 15 minutes and then after 30 minutes. However, as shown in the scatter plots from N = 7 slices on the right Fig.3c, Cal520 transients tend to increase in size over time, on average by ∼8% every five minutes. We therefore assessed another Ca^2+^ indicator (Calbryte520) that was reported to have larger retention with respect to Fluo4^43^. Calbryte520 gave transients that are similar in size to those of Cal520 and also these signals had no rundown when recordings were repeated over 30 minutes, a behaviour observed in all N = 7 slices assessed with this dye (scatter plots on Fig.3d). But in contrast to Cal520, no tendency to increase for signals with Calbryte520 was observed over time. We concluded that both Cal520 and Calbryte520 do not exhibit rundown over time, but that Calbryte520 is preferable to be combined with EF-630 for stable recordings over at least 30 minutes.

### 3.3 Combining EF-630 and Calbryte520 fluorescence recordings

Combining optimal measurements from two independent indicators requires no cross talk between the two excitation/emission channels, i.e. fluorescence signals from each indicator must be negligible when excited and recorded at the wavelengths of the other indicator. Calbryte520 has fluorescence spectra similar to fluorescein: fluorescence is not excited by red light and there is no IR emission. Consistently, in the slice of Fig.4a loaded with Calbryte520 only, Ca^2+^ transients were measured when fluorescence was excited at 470 nm and recorded at 530 ± 21 nm, but not when it was excited at 640 nm and recorded at 728 ± 64 nm, even by using >3 times laser power of what used for EF-630 imaging. In the case of EF-630, the left wing of the excitation spectrum includes blue wavelengths, but the VSD does not emit green light. Thus, in the slice of Fig.4b loaded with EF-630 only, V_m_ transients were measured when fluorescence was excited at 640 nm and recorded at 728 ± 64 nm, but not when it was excited at 470 nm and recorded at 530 ± 21 nm, even by using >3 times LED power of what used for Calbryte520 imaging.

**Fig. 4.**
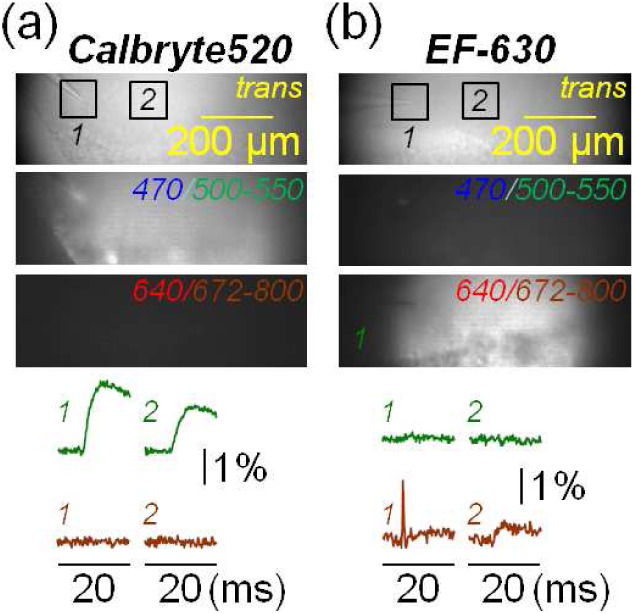
Absence of crosstalk between EF-630 and Calbryte520. (a) Top, in a slice loaded with Calbryte520 only, *trans* image and images obtained with epi-fluorescence excitation at 470 nm and emission band-pass filtered at 531±21 nm, or with epi-fluorescence excitation at 640 nm and emission band-pass filtered at 728±64 nm. Bottom, fluorescence transients from ROIs *1* and *2* illustrated above, using the two epi-fluorescence spectral windows; the Ca^2+^ transients were observed only in the green emitting window. (b) Same as a, but in a slice with EF-630 only. The V_m_ transients were observed only in the red-IR emitting window. The lookup scale of fluorescence images is the same while the 640 nm laser power was >3 times higher in the image of panel a, and the 470 nm LED power >3 times higher in the image of panel b.

It follows that EF-630 and Calbryte520 can be combined in the same experiment to obtain independent V_m_ and Ca^2+^ measurements. In the example of Fig.5a (average of 7 trials for V_m_ recordings and 2 trials for Ca^2+^ recordings), we delivered two stimuli at 50 ms interval. The spatial evolution of V_m_ and Ca^2+^ optical transients, captured from 8 instants (*I* to *VIII*) is illustrated using two distinct colour scales on the right. In the area adjacent to the electrode, the “spike” associated with direct stimulation precedes the Ca^2+^ transient, and it also precedes “EPSPs” in the CA1 region. To analyse signals, we focussed in this case to three ROIs of 50X50 pixels: ROI*1*, centred on the tip of the stimulator; ROI*2*, centred at ∼200 µm from the stimulator; and ROI*3* centred at ∼400 µm from the stimulator (Fig.5a, bottom traces). Interestingly, at ROI*1* and ROI*3*, both V_m_ and Ca^2+^ signals associated with the second pulse were larger with respect to the signals associated with the first pulse, a result which is consistent with the known phenomenon of paired-pulse facilitation^44,45,46^. To quantitatively assess signals beyond the noise, V_m_ traces in ROI*1* and all traces in ROI*2* and ROI*3* were smoothed with a Savitsky-Golay algorithm. Signals on time windows of 20 ms with first and second spike occurring after 8 ms are shown in Fig.5b. The paired-pulse ratios (p-pR) of EPSPs and Ca^2+^ transients and the delays between the spike and signals maxima (Δt) were calculated, in this way providing for each slice experiment a set of 15 quantitative parameters.

**Fig. 5.**
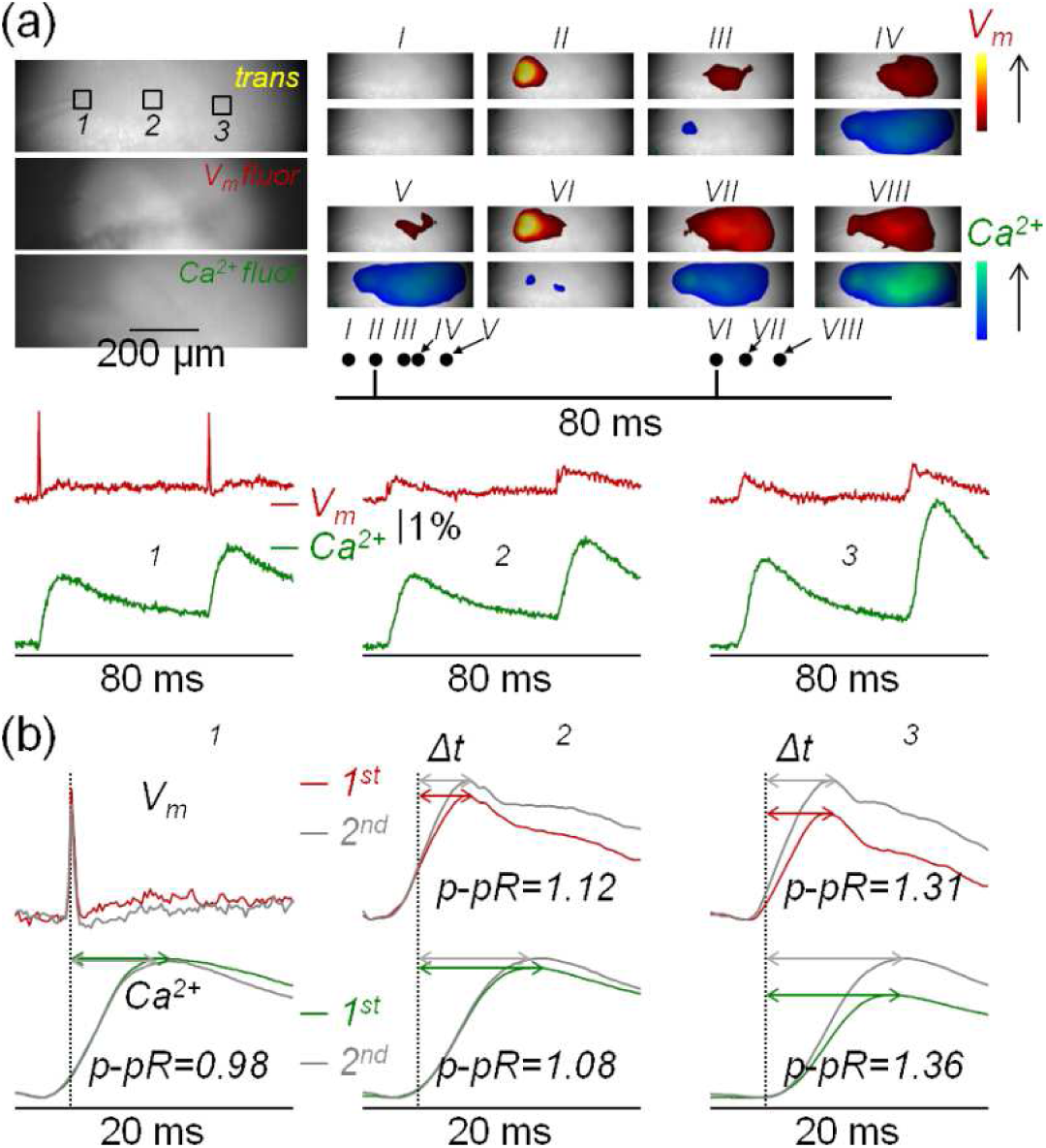
Combined EF-630 and Calbryte520 imaging. (a) Top-left, *trans*, *V_m_ fluor* and *Ca^2+^ fluor* images of the recording area with three ROIs of 50X50 pixels indicated. Top-right, spatial evolution of V_m_ and Ca^2+^ transients elicited by paired-pulse stimulation captured from 8 instants (*I* to *VIII*) illustrated using two colour scales on the right. Bottom, V_m_ (average of 7 trials) and Ca^2+^ (average of 2 trials) transients from the three ROIs; a spike is detected in ROI*1*and EPSPs are detected in ROI*2* and ROI*3*. (b) Top-left traces are spikes in ROI1 associated with the 1^st^ and 2^nd^ stimuli of the paired-protocols superimposed. The other traces are the V_m_ and Ca^2+^ transients in the ROIs indicated smoothed by a Savisky-Golay filter. The paired-pulse ratios (p-pR) of EPSPs and Ca^2+^ transients are reported and delays between the spike and signals maxima (Δt) are indicated by double-headed arrows.

Values of these parameters obtained in N = 12 slices are reported in Table 1.

**Table 1.** Values of paired-pulse ratios (p-pR) of EPSPs and Ca^2+^ transients and the delays between the spike and signals maxima (Δt) from slices where combined EF-630 and Calbrite520 imaging was performed.

| | ROI1 (stim) | | | ROI2 ~200 $\mu m$ from stim | | | | | | ROI3 ~400 $\mu m$ from stim | | | | | |
| --- | --- | --- | --- | --- | --- | --- | --- | --- | --- | --- | --- | --- | --- | --- | --- |
| | $Ca^{2+}$ | | | $Ca^{2+}$ | | | EPSP | | | $Ca^{2+}$ | | | EPSP | | |
| | p-pR | $\Delta t$ (ms) | | p-pR | $\Delta t$ (ms) | | p-pR | $\Delta t$ (ms) | | p-pR | $\Delta t$ (ms) | | p-pR | $\Delta t$ (ms) | |
| slice1 | 0.98 | 7.2 | 6.2 | 1.07 | 9.6 | 8.6 | 1.12 | 4 | 4 | 1.36 | 9.8 | 10 | 1.31 | 5.6 | 5.8 |
| slice2 | 0.95 | 6.8 | 6.2 | 1.04 | 8.6 | 8.6 | 1.19 | 3.8 | 3.8 | 1.21 | 11.4 | 9.2 | 1.15 | 5 | 4.6 |
| slice3 | 0.98 | 6.2 | 5.6 | 0.97 | 7.4 | 6.6 | 1.63 | 3.8 | 3.8 | 0.99 | 8.6 | 7.8 | 1.75 | 4.8 | 4.8 |
| slice4 | 0.96 | 6.4 | 6 | 0.93 | 9 | 8.6 | 1.10 | 5.2 | 5 | 1.01 | 11.8 | 13.0 | 1.76 | 5.4 | 5.8 |
| slice5 | 1.04 | 8.4 | 7.8 | 1.15 | 10.2 | 9.6 | 1.21 | 3.8 | 5.0 | 1.31 | 10.6 | 9.6 | 1.22 | 4 | 5.4 |
| slice6 | 1.12 | 6.2 | 5.4 | 0.93 | 8.6 | 7.2 | 1.38 | 5.6 | 5 | 1.08 | 10.2 | 9 | 1.70 | 6 | 5.2 |
| slice7 | 1.12 | 5.8 | 6.8 | 0.99 | 6.8 | 7.6 | 1.08 | 3.2 | 4 | 0.99 | 9.4 | 7.4 | 1.33 | 5.2 | 5.2 |
| slice8 | 0.94 | 6.6 | 7 | 1.29 | 9.2 | 10.8 | 1.38 | 4.4 | 5 | 1.48 | 10.6 | 9.8 | 1.5 | 5.4 | 5.6 |
| slice9 | 1.00 | 6.2 | 5.2 | 1.44 | 6.6 | 7.4 | 2.22 | 3.6 | 3.8 | 1.31 | 9.8 | 10 | 2.56 | 4.4 | 5.2 |
| slice10 | 0.99 | 7 | 6.6 | 1.35 | 9 | 9.8 | 1.39 | 4.6 | 5 | 1.53 | 10.2 | 10 | 1.5 | 5.4 | 5.6 |
| slice11 | 0.92 | 6.6 | 7 | 1.11 | 8.2 | 10.8 | 1.43 | 3.8 | 4 | 1.35 | 8.4 | 9.2 | 1.80 | 4.2 | 4.4 |
| slice12 | 0.94 | 6.6 | 6.8 | 1.01 | 8 | 8.2 | 1.8 | 3.2 | 3.4 | 1.05 | 9.4 | 9.8 | 1.84 | 3.6 | 4 |

In the directly stimulated ROI*1*, Ca^2+^ p-pR was 1.00 ± 0.06 and Δt values were 6.7 ± 0.6 ms and 6.4 ± 0.7 ms for the 1^st^ and 2^nd^ pulse respectively, suggesting that these signals are produced by Ca^2+^ influx in directly stimulated cells. V_m_ p-pR was 1.41 ± 0.32 in ROI*2* and 1.62 ± 0.36 in ROI*3*, but Ca^2+^ p-pR was only 1.11 ± 0.16 in ROI2 and 1.22 ± 0.19 in ROI3. The smaller p-pR values for Ca^2+^ with respect to V_m_ suggest that an important fraction of the Ca^2+^ transient does not have postsynaptic origin. In ROI*2*, at ∼200 µm from the electrode, V_m_ Δt values were 4.1 ± 0.7 ms and 4.3 ± 0.6 ms for the 1^st^ and 2^nd^ pulse respectively. These values were significantly shorter (P < 0.01, paired t-test) than those in ROI*3* at ∼400 µm from the electrode: 4.9 ± 0.7 ms and 5.1 ± 0.5 ms. Finally, in ROI*2*, Ca^2+^ Δt values were 8.4 ± 1.0 ms and 8.7 ± 1.3 ms for the 1^st^ and 2^nd^ pulse respectively. In ROI*3*, Ca^2+^ Δt values were 10.0 ± 1.0 ms and 9.6 ± 1.3 ms, for the 1^st^ and 2^nd^ pulse respectively. Finally, in both ROI*2* and ROI*3* and for both 1^st^ and 2^nd^ pulses, V_m_ Δt was significantly shorter than Ca^2+^ Δt (P < 0.001, paired t-test). Quantitative parameters introduced in this section were used to characterise the origin of EF-630/Calbryte520 signals.

### 3.4 Initial characterisation of the origin of EF-630/Calbryte520 signals

The final goal of this pilot study was to link experimental results to electrical and Ca^2+^ activity of the slice. The scheme of Fig.6 illustrates the sequential steps expected after stimulation.

**Fig. 6.**
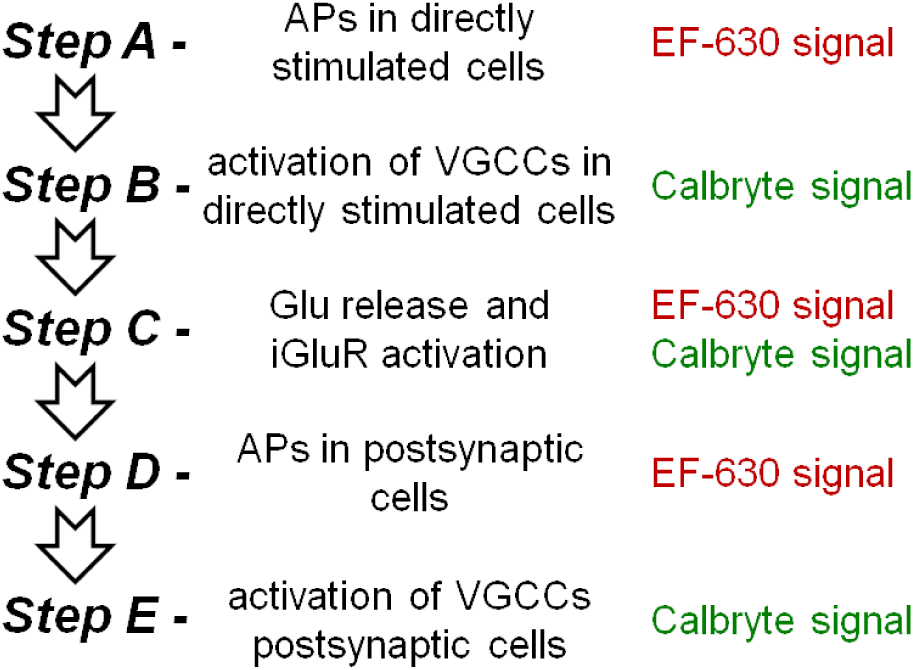
Expected sequential steps after stimulation. (*Step A*) APs in a cell population are elicited directly by the stimulation producing EF-630 signals. (*Step B*) Propagating APs activate VGCCs producing Calbryte520 signals. (*Step C*) Glutamate release activates iGluRs generating EPSPs that produce both EF-630 and Calbryte520 signals. (*Step D*) EPSPs can generate postsynaptic APs producing EF-630 signals. (*Step E*) Postsynaptic APs activate VGCCs producing Calbryte520 signals.

First (*Step A*), APs in a cell population are elicited directly by the stimulation producing EF-630 signals. These propagating APs activate voltage-gated Ca^2+^ channels (VGCCs, *Step B*) producing Calbryte520 signals and neurotransmitter release. If neurotransmitter is glutamate, this activates ionotropic glutamate receptors (iGluRs, *Step C*) and EPSPs that can be measured with EF-630, but Ca^2+^ influx via iGluRs can also produce Calbryte520 signals. Postsynaptic depolarisation above firing thresholds can boost EF-630 signals (*Step D*), activating postsynaptic VGCCs that generate Calbryte signals (*Step E*). The scheme can be repeated in further steps in case of polysynaptic activity. Since these steps are sequential, selective intervention at each step would modify the following steps leaving the previous ones unaltered. Thus, to provide an initial characterisation of EF-630/Calbryte520 signals, we blocked *Step A* (presynaptic APs) by delivering 1 µM of the Na^+^ channel inhibitor TTX, or we blocked *Step C* (iGluRs activation) by delivering a cocktail of the AMPA receptors inhibitor NBQX (10 µM) and of the NMDA receptors inhibitor D-AP5 (50 µM). As shown in the representative example of Fig.7a, TTX delivery blocks V_m_ and Ca^2+^ signals in all ROIs (described in the previous section), as expected from the scheme. In contrast, as shown in the representative example of Fig.7b, iGluRs inhibition blocks EPSPs at ROIs *2-3*, but has negligible effect on APs and Ca^2+^ transients at ROI*1*, as expected from the fact that these signals are produced at *Step A* and *Step B*. Notably, Calbryte520 transients at ROIs *2-3* were reduced but not fully blocked by NBQX and D-AP5 delivery, indicating that residual signals originate at *Step A* and *Step B* and are therefore from presynaptic axons of directly stimulated neurons. The recording of V_m_ and Ca^2+^ signals following the paired-pulse protocol before and after TTX delivery was performed in N = 5 slices. Transients were blocked in all slices (P < 0.001, paired t-test) and bar diagrams of Fig.7c show the mean ± SD of the ratios between signal maxima (maxR) after and before TTX delivery. The recording of V_m_ and Ca^2+^ signals following the paired-pulse protocol before and after delivery of iGluR inhibitors was performed in N = 7 slices and bar diagrams of the mean ± SD of maxR are shown in Fig.7d. In all slices, APs and Ca^2+^ transients in ROI1 were unaffected whereas EPSPs in ROIs *2-3* were blocked (P < 0.001, paired t-test). Yet, Ca^2+^ transients at ROIs *2-3* were reduced but not fully blocked by delivery of iGluR inhibitors.

**Fig. 7.**
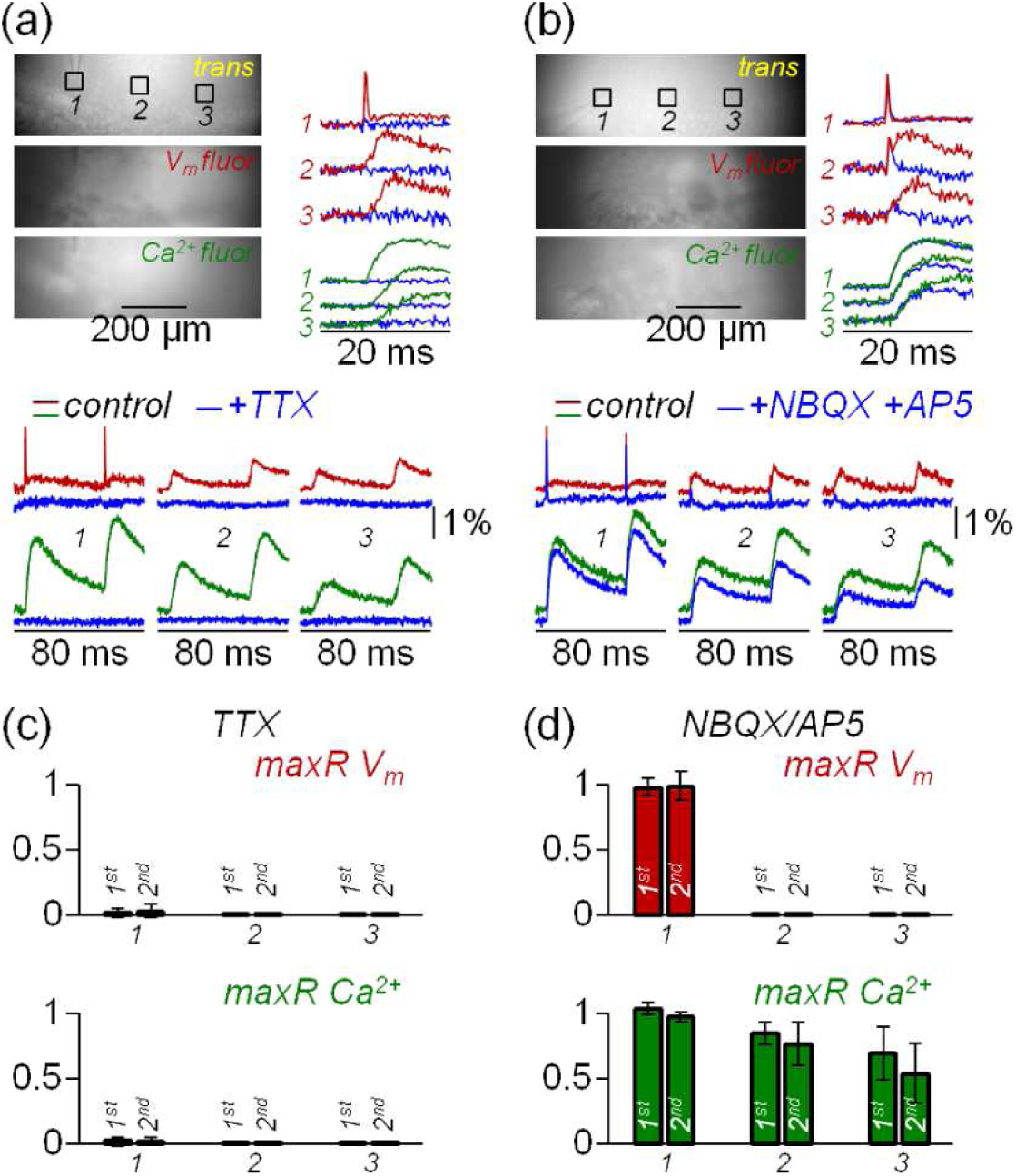
Effects of TTX and iGluR inhibitors on V_m_ and Ca^2+^ transients. (a) Top-left, *trans*, *V_m_ fluor* and *Ca^2+^ fluor* images of the recording area with three ROIs of 50X50 pixels indicated (same relative position as in Fig.5). Bottom, average V_m_ (7 trials) and Ca^2+^ (2 trials) transients elicited by paired-pulse stimulation from the three ROIs under control conditions and after delivery of 1 µM TTX (blue traces). Top-right, V_m_ and Ca^2+^ transients associated with the 1^st^ stimulation pulse, in control and after TT delivery, are superimposed. (b) Same as panel a, but in an example of delivery of 10 µM NBQX and 50 µM D-AP5 (iGluR inhibitors). (c) Mean ± SD (N = 7 slices) of the ratios between signal maxima (maxR) after and before TTX delivery: top, V_m_ transients; bottom, Ca^2+^ transients. (d) Same as in panel c, but for N = 7 slices where delivery of iGluR inhibitors was tested.

Experiments of Fig.7b and Fig.7d were used to evaluate the quantitative parameters related to Ca^2+^ transients reported in Table 1. Ca^2+^ transients of Fig.7b, smoothed with a Savitsky-Golay filter, are shown in Fig.8a. Specifically, as Ca^2+^ transients associated with the 1^st^ and 2^nd^ stimulation pulse were superimposed, it was evident in ROI*2* and ROI*3*, but less in ROI*1*, that delivery of iGluR inhibitors decreased the p-pR and the delay between spike and Ca^2+^ signal peaks (Δt). Scatter plots in Fig.8b show the ratios of p-pR (p-pRR) and of Δt (ΔtR) parameters in the 7 slices reported in Fig.7d. Delivery of NBQX and D-AP5 systematically reduced p-pR and Δt of Ca^2+^ transients in both ROI*2* and ROI*3* (P < 0.01 paired t-test). The first result indicates that Ca^2+^ paired-pulse facilitation in control conditions is mediated by glutamatergic synaptic transmission. The second result gives a quantitative estimate between the presynaptic and postsynaptic components of Ca^2+^ transients.

**Fig. 8.**
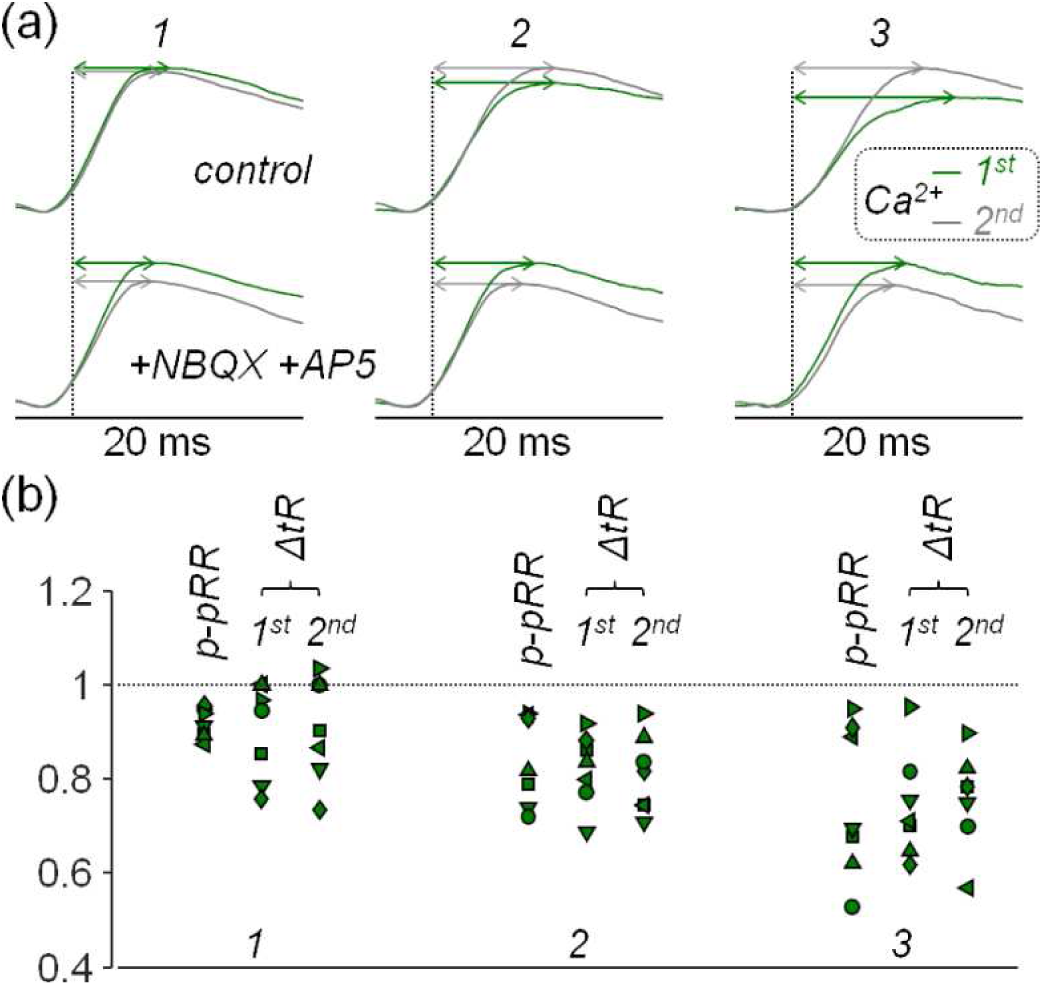
Effects iGluR inhibitors on Ca^2+^ p-pR and Δt parameters. (a) Ca^2+^ transients of Fig.7b in the three ROIs indicated smoothed by a Savisky-Golay filter; the transients associated with the 1^st^ and 2^nd^ stimulating pulses are superimposed; delays between the spike and signals maxima (Δt) are indicated by double-headed arrows; top traces are in control conditions; bottom traces are after delivery of iGluR inhibitors. (b) Scatter plots in the three ROIs of the ratios between p-pR (p-pRR) and Δt (ΔtR) parameters after delivery of NBQX and D-AP5 and in control conditions from N = 7 slices where iGluR inhibitors were tested.

## 4 Discussion

In the present article we report substantial improvements in brain slice V_m_ and Ca^2+^ imaging achieved using the combination of the VSD EF-630 and of the Ca^2+^ indicator Calbryte520. The brain slice is an *ex-vivo* preparation that allows investigation of neurons in their physiological environment where many connections that form local networks are preserved. At present, it is possible to investigate brain slice activity in specific cell types using genetically encoded fluorescence indicators both for V_m_ (GEVI) ^47,48,49^ and Ca^2+^ (GECI)^50,51^. These state-of-the-art approaches achieve relatively large (>1%) fluorescence transients, but they encounter several technical challenges starting from the need of expressing the protein in a living animal, but also caveats related to the kinetics of indicators. For example, a remarkable temporal discrepancy between GEVI and GECI imaging was reported in brain slices^52^. Organic indicators, in contrast, do not offer the possibility to target cell classes with specific molecular identity, but the possibility to record fluorescence transients with size comparable to those from GEVIs or GECIs can make their use the preferable choice in many instances. Slices can be prepared in standard ways from any animal model and only later loaded with the Ca^2+^ indicator, and rapidly stained with the VSD just before recording. State-of-the-art CMOS cameras can be used to record fluorescence at high spatiotemporal resolution with stable signals lasting for >30 minutes. In addition, the large gap between fluorescence spectra of EF-630 and Calbryte520 allows recordings of fully independent V_m_ and Ca^2+^ signals associated with the same neuronal activity. Notably, the combination of blue/red excitation still leaves the possibility of a third independent illumination pathway in the UV range that can be used, for instance, for uncaging techniques^53^. Alternatively, by giving up Ca^2+^ imaging, blue illumination can be used for channelrhodopsin excitation in combination with V_m_ imaging^54^. The outstanding SNR of the measurements obtained at very high spatiotemporal resolution that are reported in this article allowed an initial characterisation of signals elicited by electrical stimulation of the hippocampal CA3 region. Clear APs from the population of directly stimulated hippocampal neurons are distinguishable, as well as monosynaptic EPSPs in the CA1 region where Shaffer collaterals are expected to make connections. These EPSPs were fully blocked by delivery of iGluR inhibitors (Fig.6b and Fig.6d). A straightforward correlation between V_m_ and Ca^2+^ signals, however, is not possible since Ca^2+^ signals at the EPSP sites comprise a combination of postsynaptic and presynaptic Ca^2+^ influx. This result is not surprising because EF-630 staining and Calbryte520 loading are independent and therefore fluorescence transients do not, in general, originate from the same cells (or if they do, not in the same proportion). Nevertheless, the accurate analysis of signal kinetics permitted by the high temporal resolution (5 kHz) allowed identifying different components of the Ca^2+^ transients and elucidating the observation of paired-pulse facilitation in postsynaptic cells. Full characterisation will require more pharmacological analysis that may include delivery of D-AP5 alone to reveal Ca^2+^ influx via NMDA receptors, blockade of VGCCs, or blockade of K^+^ channels that may result in the AP waveform changes. Such pharmacological investigations, however, are beyond the scope of the present technical article aimed at reporting a methodological step-forward in combined V_m_ and Ca^2+^ imaging ex vivo.

## 5 Conclusions

Achievements reported in this article open the door to a variety of potential studies in different brain regions (hippocampus, neocortex, thalamus, cerebellum, spinal cord, etc.), as well as in animal models of disease (epilepsy, autism, ataxia, migraine, etc.). They also lift previous limitations in detailed drug investigations of the V_m_ – Ca^2+^ signalling.

## Data and Code Availability

Frame sequences that were quantitatively analysed, either single trial of averages of 2 or 7 trials as indicated in the text, are available in the public repository Zenodo (doi: 10.5281/zenodo.17725316). For data analysis, standard Matlab or Python functions were used and Matlab scripts were written only to fasten data handling..

## Acknowledgments

This work was supported by the *Agence Nationale de la Recherche* through two grants (ANR-21-CE180042, Nav12RESCUE and ANR-11-LABX-0015, Labex *Ion Channels Science and Therapeutics*). We thank Cindy Tellier and Hervé Dubouchaud for handling mice at the animal facility.

